# A single-dose recombinant rVSVΔG-CCHFV-GPC vaccine protects nonhuman primates against heterologous Crimean-Congo hemorrhagic fever virus challenge

**DOI:** 10.64898/2026.10.21.760752

**Authors:** Robert W. Cross, Kevin Melody, Rachel O’Toole, Viktoriya Borisevich, Daniel J. Deer, Joan B. Geisbert, Courtney B. Woolsey, Krystle N. Agans, Abhishek Prasad, Dafna Abelson, Xhevat Jakupi, Lindita Ajazaj-Berisha, Salih Ahmeti, Thomas W. Geisbert

**Affiliations:** Galveston National Laboratory, University of Texas Medical Branch, Galveston, TX 77555 USA; Department of Microbiology and Immunology, University of Texas Medical Branch, Galveston, TX 77555 USA; MAPP Biopharmaceuticals, San Diego, CA, USA; Department of Microbiology, National Institute for Public Health of Kosova; University of Prishtina “Hasan Prishtina”, University Clinical Center of Kosova, Clinic of Infectious Diseases

## Abstract

Crimean-Congo hemorrhagic fever virus (CCHFV) is a widespread tickborne virus with potential to cause lethal disease in humans. Countermeasure research for CCHFV has been hindered by the lack of a preclinical model that closely recapitulates human disease in an immunocompetent species. Recent reports of experimental CCHFV infection of cynomolgus macaques indicate these animals exhibit disease reflective of that observed in humans. Here, we assessed the ability of a replication competent, recombinant vesicular stomatitis virus (rVSV) vector encoding CCHFV structural glycoproteins (rVSV-CCHFV-GPC) from a Clade III isolate to prevent disease in cynomolgus macaques after challenge with a Clade V isolate of CCHFV and demonstrate that a single vaccination dose is potently immunogenic and reduced viremia and inflammatory biomarker circulation when compared with unvaccinated controls. Importantly, humoral responses to CCHFV glycoproteins in vaccinated macaques were similar to those observed in human CCHF survivors. This study further supports that a rVSV expressing only the CCHFV-GPC has the potential to serve as a vaccine platform against CCHFV infections.

## INTRODUCTION

Potential for devastating disease, expanding geographical distribution of the reservoir, and a lack of globally approved prophylaxes have led to Crimean-Congo hemorrhagic fever virus (CCHFV) being classified as a priority pathogen by the World Health Organization^1^. CCHFV is a tick-borne, negative-sense RNA virus of the *Nairoviridae* family. The virus has a tri-segmented genome divided into S, M, and L coding regions, which encode the nucleoprotein (NP), glycoprotein (Gn/Gc) and several nonstructural proteins (ex. GP38 GP85, GP160) and polymerase^2^. The main reservoir for CCHFV are the *Hyalomma* genus of ticks, which are widely distributed across Africa, Southern and Eastern Europe, the Middle East, and Asia^3^. This expansive geographic range likely contributed to evolution into at least six distinct phylogenetic clades with 20-31% divergence between the three genomic coding segments across clades^4^. Humans are exposed to CCHFV through tick bites or direct contact with contaminated blood products. While most infected animals are not known to develop overt disease^5^, humans may develop a nonspecific febrile illness that can progress to acute hemorrhagic disease typified by progressive coagulopathy and vascular leakage sometimes leading to fatal systemic shock^6^. The global case fatality rate is approximately 30%, but outbreak size and disease severity show large geographic variablity^4^, which is likely affected by both intrinsic (e.g., virus clade, host genetic variation) and extrinsic (e.g., clinical reporting, surveillance) factors. Globalization and climate change are thought to be contributing to *Hyalomma* spp. home range expansion, which is likely increasing risk of CCHFV infections^7^. There are no widely approved vaccines against CCHFV. The only vaccine approved for use in humans is an inactivated, aluminum-adjuvanted injection based on a virus strain passaged in suckling mouse brains. While immunogenic, the regimen requires a series of three injections followed by periodic boosts and is only available in Bulgaria^8^. Multiple doses and lifetime boosts are not ideal in outbreak settings where protection is needed quickly. Additionally, vaccine compliance is likely to be increased with fewer doses.

Progress in the development of vaccines for CCHFV infection has been hindered the need for high containment facilities (i.e., Biosafety Level 3 or 4) to test the virus and the absence animal models that accurately recapitulates human disease. Outbred mice are resistant to disease^3^, but mouse strains deficient in innate immune signaling are particularly susceptible to infection and even lethal disease^9–11^. These immunodeficient mice have been employed to assess many vaccine platforms^12^; however, these models may not reflect the full potential to induce immunity in vaccinated animals due to the deletion of key immune components that facilitate susceptibility to CCHFV.

Recently, an immunocompetent nonhuman primate (NHP) model for CCHFV infection was described by multiple groups^13–15^. NHPs serve as a gold-standard for disease and therapeutic research as they are the species most genetically related to humans and tend to recapitulate disease more accurately than rodents. Cynomolgus macaques challenged intravenously (IV) with CCHFV strain Kosovo Hoti displayed lethargy, edema, epistaxis, coagulopathy, and death but severe disease was not consistent in all subjects^14^. Subsequent attempts to replicate this outcome did not induce lethal disease with either Hoti or Afghanistan strains^13,15^. Nonetheless, infected macaques became viremic and showed relevant signs of disease observed in humans such as decreased appetite, epistaxis, edema, hemorrhagic rashes, leukopenia, lymphopenia, thrombocytopenia, and elevated liver enzymes across all studies^13–15^. To date, a DNA vaccine requiring three doses is the only strategy tested for CCHFV prevention in the macaque model^16^; however, no DNA based vaccines have successfully completed all the steps necessary for licensure in humans.

Previous studies have demonstrated that rVSV-based vaccines with the glycoproteins of other high-risk pathogens elicit robust protection in NHP models of filoviruses, Lassa virus, and Nipah virus^17–26^. Such vaccine candidates are ideal because they provide strong durable immunogenicity with a single dose, have shown significant protection in as little as one-week post-dosing, and targeted immunity is not affected by previous vaccination with rVSV vectors^17,19,21,23,26,27^. The rVSV vaccine, ERVEBO^®^, was the first widely approved prophylaxis for Ebola virus disease in humans, as it demonstrated a strong safety profile, immunogenicity, and protection in clinical trials^28–31^. This licensure emphasizes the utility of rVSV vectors as a vaccine platform and suggests its potential for use against other viruses. Previously we demonstrated the protective efficacy of a recombinant vesicular stomatitis virus (rVSV) vector encoding CCHFV structural glycoproteins (rVSV-CCHFV-GPC) to protect 100% of vaccinated immunodeficient mice from lethal infection with the Turkey2004 strain of CCHFV^32^. In this work, cynomolgus macaques were vaccinated with a single dose of VSV-CCHFV-GPC then challenged with the Hoti strain of CCHF and evaluated for protection from disease.

## RESULTS

### Vaccination of cynomolgus macaques reduces viremia and disease

To test the efficacy of VSV-CCHFV-GPC in a higher order model, we vaccinated five macaques with a single intramuscular (IM) dose. The vector was well tolerated, and no adverse events were observed post-vaccination (data not shown). Thirty-five days post-vaccination, we challenged the vaccinated group along with three unvaccinated control macaques with Hoti by intravenous (IV) injection. Viral RNA (vRNA) was detected in all subjects where lower mean levels were recorded in vaccinated through the study (Fig 1A) and overall peak vRNA levels were higher in unvaccinated animals (Figure 1B; Mann Whitney (p=0.0357)). vRNA was cleared to undetectable levels faster in unvaccinated animals compared to unvaccinated controls (Figure 1C, Mann Whitney (p=0.0357)). Circulating infectious virus was detected in all subjects up to day 3 post challenge and lower mean titers were recorded in vaccinated animals up to this point post challenge (Fig 1D). Mean peak infectious titers were highest in unvaccinated animals (Figure 1E; Mann Whitney (p=0.0357)). Infectious virus was cleared in vaccinated animals up to 4 days faster in vaccinated animals compared to unvaccinated controls (Figure 1F). At study endpoint, vRNA in a panel of tissues were assayed. Except for one vaccinated animal which had no detectable vRNA in any tissue, only immune organs (axillary lymph nodes, inguinal lymph nodes, and spleen) were positive for vRNA with no significant differences for any tissue despite vaccination status (Supplementary Table 1).

**Figure 1:**
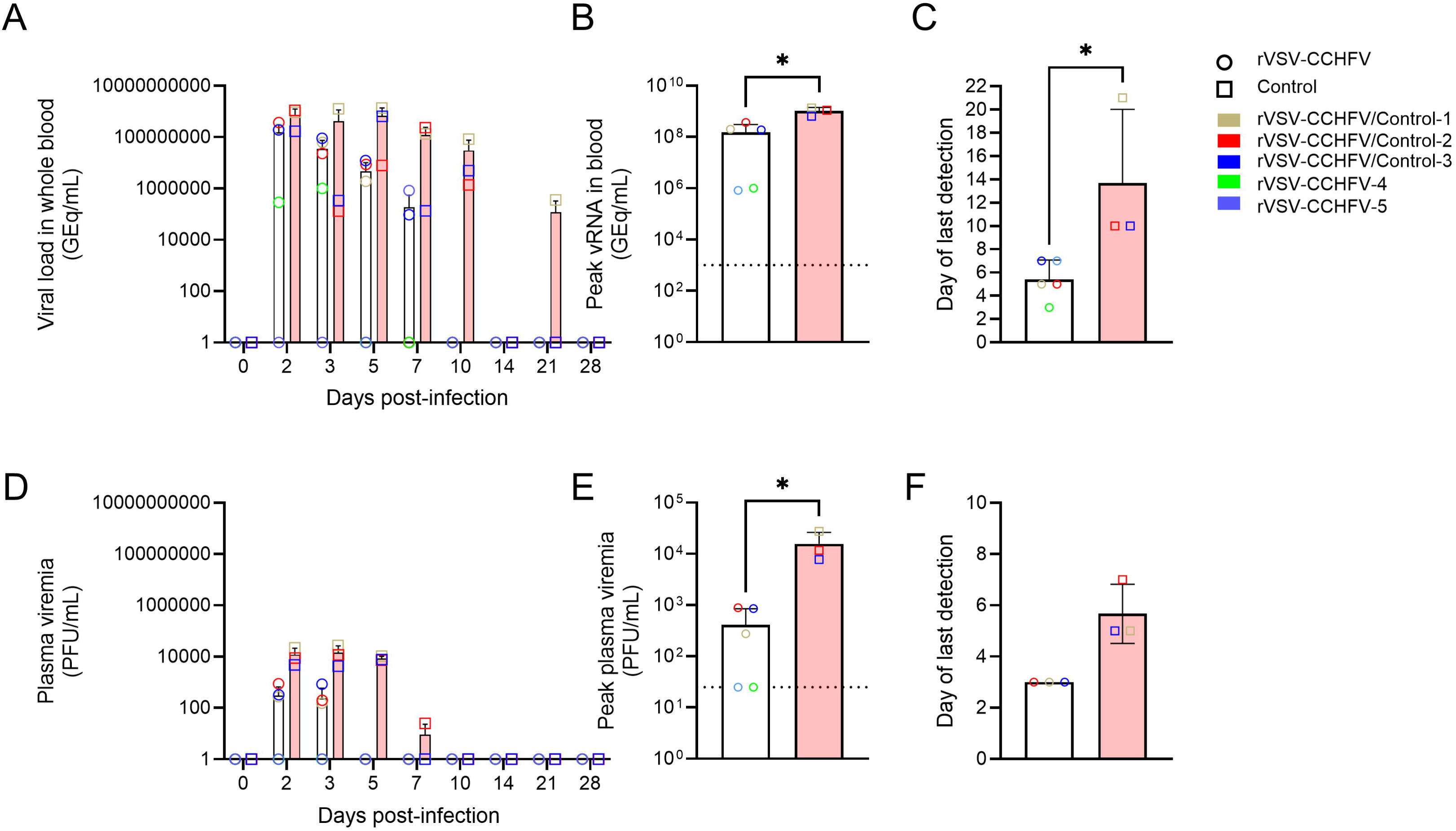
Viremia in vaccinated and unvaccinated CCHFV-infected macaques. Concentrations of vRNA in whole blood (A), Peak vRNA (B), and day of last vRNA detection (C), infectious virus in plasma (D) Peak infectious virus (E), and day of last infectious virus detection (F) were detected by RT-qPCR and plaque assay, respectively. Mann-Whitney tests were utilized to compare means where * indicates a p value ≤ 0.05.

In line with our previous data^13^, clinical disease was mild in all control subjects with no lethal outcome. No subject met clinical scoring criteria, and none developed fever or weight fluctuations greater than 10% from baseline (data not shown). Only the unvaccinated subjects exhibited decreased appetite during the acute phase of disease (Supplementary Fig. 1A).

All subjects displayed changes in hematology and serum chemistry parameters, but the vaccinated group displayed fewer overall changes (Table 1). Neutropenia and basopenia were noted in all subjects; however, only fold-changes in red blood cell counts, While all serum chemistry parameters were modestly affected post-infection with higher mean values were recorded in unvaccinated animals suggestive of greater pathological insult than observed in vaccinated animals (Supplemental Fig. 1B, 1C), otherwise most chemistry parameters did not fluctuate appreciably nor did any display significant difference between groups. By day 28, all hematological and chemistry parameters were near pre-infection levels in all subjects.

**Table 1.** Clinical description of vaccinated and unvaccinated CCHFV-infected cynomolgus macaques following CCHFV Hoti challenge.

| Treatment | Subject No. | Sex | Clinical illness<sup>a, b</sup> | Clinical pathology<sup>b, c</sup> |
| --- | --- | --- | --- | --- |
| Not vaccinated | CONTROL 1 | F | Decreased appetite (d1, 4-8) | Leukopenia (d2-7, 21, 28); thrombocytopenia (d5-7); monocytopenia (d2-5, 10-28); neutropenia (d2-28); eosinopenia (d2, 5-7, d8); basopenia (d2, 5-7, 28); lymphocytosis (d10-21); ↓GLU (d28); ↓BUN (d5); ↓CRE (d3, 14); ↑AST (d2-7); ↑CRP (d2); ↓CRP (d5, 14) |
|  | CONTROL 2 | F | Decreased appetite (d4-8, 10-11, 16) | Leukopenia (d2-7); thrombocytopenia (d2-7); lymphopenia (d2, 4); monocytopenia (d2-5, 28); neutropenia (d2-21); eosinopenia (d2-7); basopenia (d2-7); lymphocytosis (d10-21); monocytosis (d7-10); eosinophilia (d10); ↓CRE (d2, 5-7, 14, 28); ↑ALT (d2-3); ↑AST (d2-7); ↑AMY (d10, 28); ↑CRP (d2, 28) |
|  | CONTROL 3 | M | Decreased appetite (d5) | Leukopenia (d2, 5); thrombocytopenia (d5-7); lymphopenia (d5); neutropenia (d2, 5, 10-28); eosinopenia (d2, 5); basopenia (d2, 5); lymphocytosis (d10-14); monocytosis (d10-14); eosinophilia (d7-28); basophilia (d10-21); ↑AST (d3-7); ↓ALP (d10); ↓AMY (d2-5, 10); ↑CRP (d2) |
| Vaccinated | rVSV-CCHFV-1 | M | None | Leukopenia (d2, 5); lymphopenia (d2); neutropenia (d2, 5-7); eosinopenia (d2-5); basopenia (d2-5); monocytosis (d2-3, 7, 14-21); eosinophilia (d21); ↑AST (d3-5); ↑CRP (d2) |
|  | rVSV-CCHFV-2 | M | None | Thrombocytopenia (d5); neutropenia (d5-10, 21-28); basopenia (d5); monocytosis (d3-10); eosinophilia (d2-3, 10-28); basophilia (d10-14); basophilia (d10); ↓CRE (d5, 10); ↑AST (d3-5); ↑CRP (d2-3) |
|  | rVSV-CCHFV-3 | M | None | Leukopenia (d2, 28); thrombocytopenia (d2-5); lymphopenia (d2); monocytopenia (d2, 28); neutropenia (d2, 5-7, 14-28); eosinopenia (d2); basopenia (d2, 5); lymphocytosis (d7-14); monocytosis (d7); eosinophilia (d28); ↓CRE (d2, 5-28); ↑AST (d3-7); ↓AMY (d2); ↑CRP (d2-3) |
|  | rVSV-CCHFV-4 | F | None | Lymphopenia (d2); neutropenia (d5, 28); eosinopenia (d2, 5, 28); basopenia (d2, 5, 28); monocytosis (d2-7, 14, 28); neutrophilia (d3); ↓AST (d14); ↑CRP (d2) |
|  | rVSV-CCHFV-5 | F | None | Monocytopenia (d28); neutropenia (d2-3, 10-28); basopenia (d2); ↓CRE (14); ↑AST (d3); ↑CRP (d2) |

<sup>a</sup> values in parentheses represent days post-challenge.
<sup>b</sup> decreased appetite is defined as ≥25% decrease in daily feed intake. Anorexia is defined as no consumption of daily feed.
<sup>c</sup> Leukopenia, thrombocytopenia, lymphopenia, monocytopenia, neutropenia, eosinopenia, and basopenia are defined as a ≥35% decrease from baseline in numbers of leukocytes, platelets, lymphocytes, monocytes, neutrophils, eosinophils, and basophils, respectively. Leukocytosis, thrombocytosis, lymphocytosis, monocytosis, neutrophilia, eosinophilia, and basophilia are defined as 2-fold increase from baseline in numbers of leukocytes, platelets, lymphocytes, monocytes, neutrophils, eosinophils, and basophils, respectively.
<sup>d</sup> ↓ represents a ≥25% decrease and ↑ represents a ≥2-fold increase in blood chemistry parameters relative to baseline values.
ALB, albumin; ALT, alanine aminotransferase; ALP, alkaline phosphatase; AMY, amylase; AST, aspartate aminotransferase; BUN, blood urea nitrogen; CA, calcium; CRE, creatinine; CRP, C-reactive protein; GGT, gamma glutamyltransferase; GLU, glucose; TP, total protein; UA, uric acid

**Table 2.** Detection by q-RT-PCR of vaccinated and unvaccinated CCHFV-infected cynomolgus macaques RNA in tissue samples collected at study endpoint, 28 days post-challenge.

| Treatment | Subject No. | Axillary lymph node | Inguinal lymph node | Spleen | Liver | Lung | Pancreas | Adrenal gland | Bladder | Brain<sup>b</sup> | Reproductive tissues<sup>c</sup> | Conjunctiva/ Eye |
| --- | --- | --- | --- | --- | --- | --- | --- | --- | --- | --- | --- | --- |
| Not vaccinated | CONTROL 1 | —<sup>a</sup> | 7.85 | 7.07 | - | - | - | - | - | - | - | - |
|  | CONTROL 2 | 7.54 | 7.95 | 5.22 | - | - | - | - | - | - | - | - |
|  | CONTROL 3 | 7.47 | 7.81 | - | - | - | - | - | - | - | - | - |
| Vaccinated | rVSV-CCHFV-1 | - | 7.24 | 5.15 | - | - | - | - | - | - | - | - |
|  | rVSV-CCHFV-2 | 4.43 | 5.49 | - | - | - | - | - | - | - | - | - |
|  | rVSV-CCHFV-3 | - | - | - | - | - | - | - | - | - | - | - |
|  | rVSV-CCHFV-4 | - | - | - | - | - | - | - | - | - | - | - |
|  | rVSV-CCHFV-5 | - | 7.85 | 7.07 | - | - | - | - | - | - | - | - |
| <sup>a</sup> values representative of log<sub>10</sub> GEq/ml. — indicates data below limit of detection, 3.00 log<sub>10</sub> GEq/ml<br><sup>b</sup> frontal cortex, brain stem, and cerebral spinal cord analyzed separately<br><sup>c</sup> gonads and uterus or prostate analyzed separately |  |  |  |  |  |  |  |  |  |  |  |  |

To further investigate differences between groups, we measured a panel of circulating serum cytokines previously demonstrated to be affected by CCHFV infection in macaques^13^. Reduced amounts of circulating Transforming growth factor alpha (TGF-α), G-CSF (Granulocyte Colony-Stimulating Factor), Interferon gamma (IFN-g), Interleukin 10 (IL-10), and Interleukin 1b (IL-1b) were noted in vaccinated animals suggestive of a dampened proinflammatory response compared with unvaccinated animals (Fig. 2). Because CCHFV infection can lead to varied coagulopathies in humans^33–36^, we analyzed changes in fibrinolysis- and thrombosis-related proteins during the acute phase of infection. While there were no statistical differences observed, increased mean levels Plasminogen activator inhibitor-1 (PAI-1), D-Dimers, and Tissue Plasminogen Activator (tPA) suggestive of a more pronounced activation of coagulation pathways in unvaccinated animals was observed (Fig. 3).

**Figure 2.**
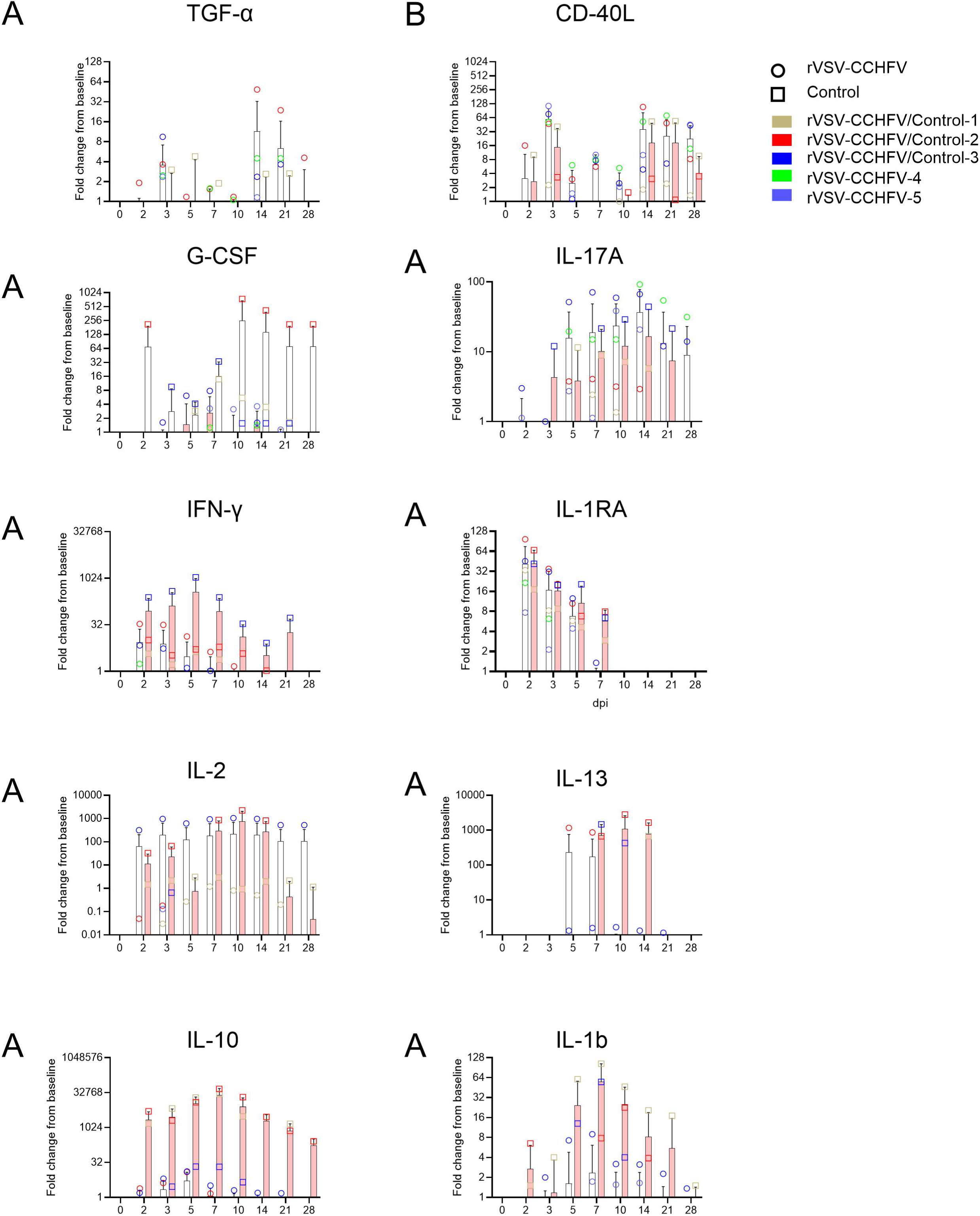
Changes in serum cytokine concentrations in vaccinated and unvaccinated CCHFV-infected cynomolgus macaques. Serum concentrations of TGF-A (A), CD40L (B), G-CSF (C), IL-17A (D), IFN-G (E), IL-1RA (F), IL-2 (G), IL-13 (H), IL-10 (I), AND IL-1B (G) were measured and normalized to pre-infection values. Dashed line represents pre-challenge baseline. Each data point is the mean of two technical replicates. Bars represent mean and standard deviation of groups.

**Figure 3.**
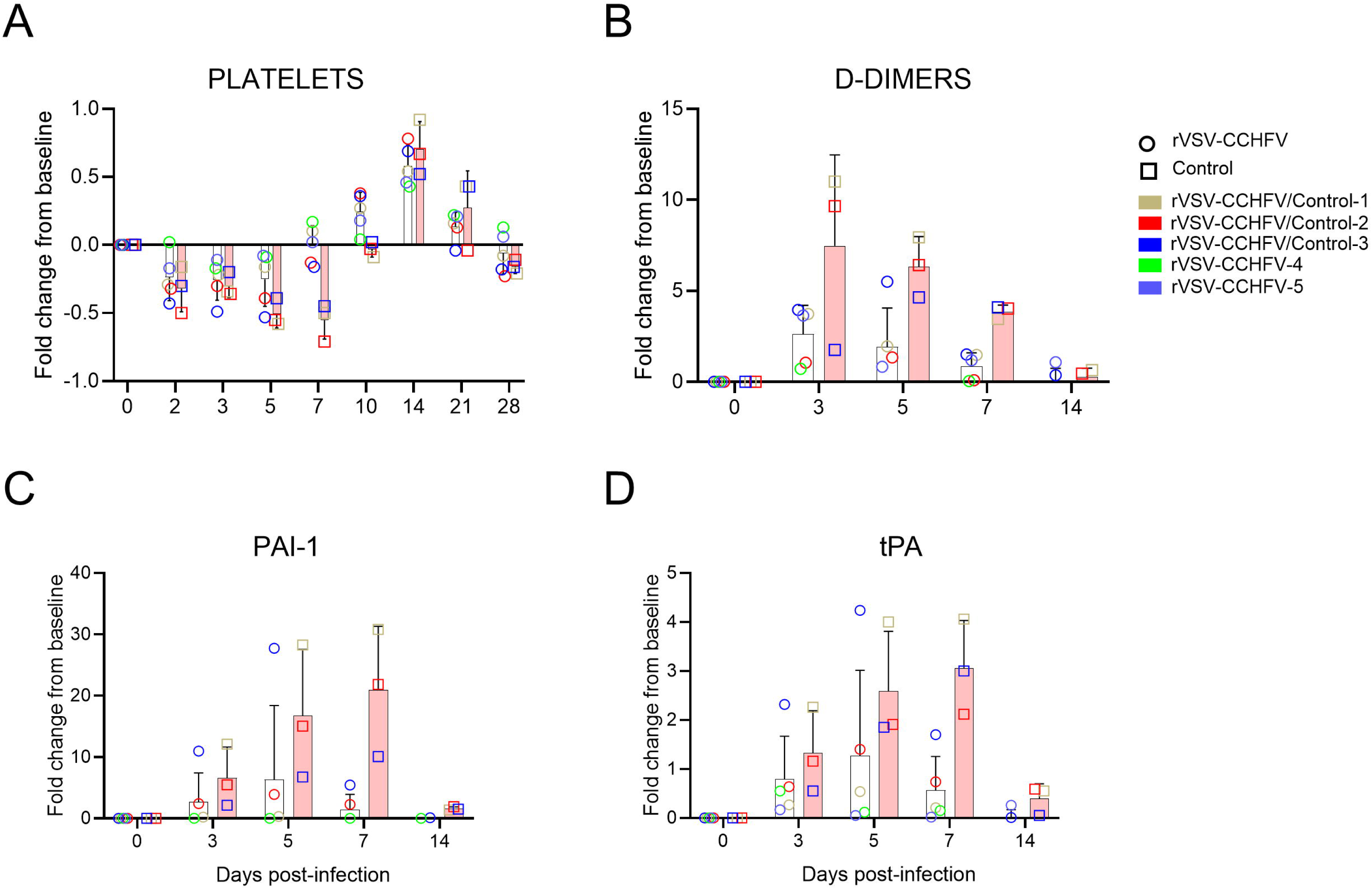
Changes in plasma thrombosis-related proteins in vaccinated and unvaccinated CCHFV-infected cynomolgus macaques. Fold-changes circulating thrombocytes (A), in serum concentrations of D-dimers (B), PAI-1 (C), and tPA (D) were measured with using a fluorescence-based bead assay. Each data point is the mean of two technical replicates. Bars represent mean and standard deviation of groups.

### Vaccination elicits an anti-CCHFV antibody response

To determine immunogenicity of the VSV-CCHFV-GPC vaccine, we measured serum antibody levels against CCHFV NP, Gc, or GP38. The unvaccinated group developed higher anti-NP IgG responses (Fig. 4A); which is in line with natural recovery observed in the human response. Because the rVSV vector did not express CCHFV NP, we measured antibody levels to recombinant CCHFV M products. The vaccinated group developed an earlier and overall higher anti-Gc response than the control group (Fig. 4B). The vaccinated group also developed an earlier and stronger response to the non-structural M product, GP38 (Fig. 4C). By day 28, the control group had mean anti-Gn/Gc and GP38 titers equivalent to the vaccinated group at day 5. Three VSV-CCHFV-GPC subjects had detectable post-vaccination neutralization titers at time of CCHFV challenge and all subjects, regardless of vaccination status, developed a neutralization response by the study endpoint (Fig. 4D).

**Figure 4.**
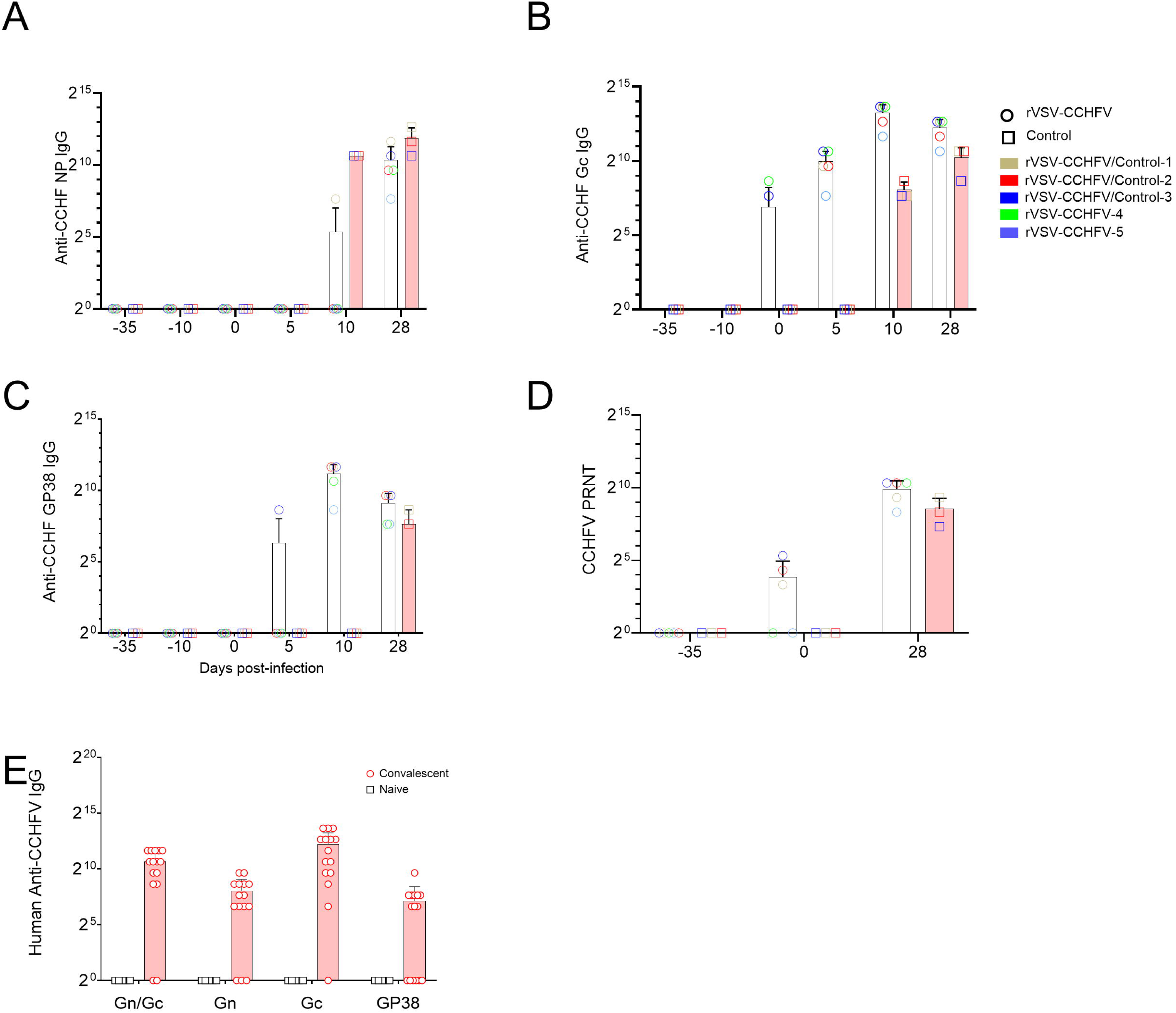
Humoral antibody response in vaccinated and unvaccinated CCHFV-infected cynomolgus macaques and convalescent humans exposed to CCHFV. IgG response was measured against CCHFV NP (A), purified Gc (B) or purified GP38 (C). Serum 50% plaque reduction neutralization values were also measured (D). Each data point is the mean of two technical replicates. Dashed lines designate the limit of detection for each assay. Antibody responses from convalescent serum collected from individuals who were positive/hospitalized for CCHFV infection (n = 15) and uninfected controls (n = 5) were measured by ELISA. Each data point is the mean of two technical replicates. Bars represent mean and standard deviation of groups. Dashed line designates the limit of detection for the assays.

### Anti-CCHFV glycoprotein antibody response from convalescent humans

To determine if the antibody response we observed in both vaccinated and control animals mirrors natural infection in humans, we measured the anti-CCHFV IgG responses of convalescent human serum from 15 confirmed survivors from Kosovo and five CCHFV-naïve persons using a suite of recombinant CCHFV antigens. We detected antibody responses to both Gn/Gc and GP38 in humans previously infected by CCHFV; though, no sample was uniformly positive to all three antigens tested (Fig. 4E). Consistent with the VSV-CCHFV-GPC-vaccinated group, we observed reactivity to GP38, Gc and Gn where Gc appears to be the more immunoreactive glycoprotein epitope in humans that have survived natural infection highlighting its importance in the immune response to CCHFV and as a vaccine component.

## DISCUSSION

Given the potential for severe disease and lack of widely approved anti-CCHFV prophylaxis, we utilized the recently described cynomolgus macaque model to assess the protective efficacy of the VSV-CCHFV-GPC vaccine^32^. While natural correlates of protection against CCHFV incompletely understood, lower viral loads^34,39–42^ and a CCHFV antigen targeted humoral immune response are associated with improved clinical outcomes in humans^34,40,43,44^. VSV-CCHFV-GPC tested here satisfied both conditions as it proved to be safe, immunogenic, and protective in the vaccinated group, which developed an earlier anti-CCHFV antibody response, had lower virus titers, displayed more rapid clearance of virus and some biomarkers were less affected compared to the unvaccinated group.

Despite observing lower viral loads and more rapid clearance of both vRNA and infectious virus in the vaccinated group, disease was mild with little observable clinical differences between the two groups. Although fold-changes in blood cell counts in all subjects were transient and effects in some cases were low, we did observe a significantly reduced effect on red blood cells, thrombocytes, and hemoglobin in the vaccinated group towards the end of the acute phase of disease. Nonetheless, these biomarkers have been associated with poorer prognosis in humans^45^.

Because CCHFV infection is linked to coagulopathies, we also measured factors related to thrombosis and fibrinolysis. D-dimers, PAI-1, and tPA all displayed notable increases post-challenge in both groups suggesting perturbation of normal coagulation parameters, but all three parameters rapidly returned to baseline faster in the vaccinated group, suggesting vaccination can impact resolution of virus instigated coagulopathy. Lower concentrations of all three parameters are also associated with survival in human infections^45,46^. Because both the vaccinated and unvaccinated groups became infected with CCHFV, nearly all measured biomarkers showed acute phase fluctuations without significant differences between groups. In a three dose DNA-based vaccine study, macaques were protected from detectable viremia and disease with demonstratable differences in biomarkers, such as albumin and aspartate aminotransferase, between vaccinated and unvaccinated groups^16^. It is possible that a prime-boost strategy with VSV-CCHFV-GPC may have elicited a stronger humoral response or even sterilizing immunity, but the strength of the rVSV system is in its ability to elicit quick protection in a single dose^27^. However, CCHFV infection in macaques trends toward mild disease and a consistent severe or lethal model of infection is needed to better define correlates of disease and protection. Until such a consistent model of CCHFV infection is developed, viremia and circulating biomarkers should continue to be monitored as correlates of disease, similar to other non-lethal NHP vaccine models^47–50^.

To broaden the potential immune response of VSV-CCHFV-GPC we included the entire CCHFV M coding region, which likely allows for natural Gn/Gc processing and production of nonstructural proteins, particularly GP38. In this study, the vaccinated group developed earlier and greater humoral immune responses to CCHFV GPC products; however, only antibody titers to Gc were readily observed in the vaccinated group at time of challenge suggesting protection developed from a rapidly mounted anamnestic immune response may be important. As expected, vaccinated subjects developed high antibody binding titers targeting Gc initially, but GP38 IgG antibodies were observed by day 5. In line with previous reports, we confirmed higher binding antibodies to Gc over Gn in human survivors. This may be attributed to the larger size of Gc relative to Gn^51^ or folding and presentation differences in the purified proteins. The greater response to Gc does not diminish the role of Gn in protection as monoclonal antibodies to both antigens have shown protection in mice^37^ and their exact roles in protection, particularly in humans, requires further exploration. The antibody response to GP38 is promising as we show this secreted protein of unknown function is immunogenic in a higher order preclinical model and may also be contributing to protection. Antibodies targeting GP38 have shown protection in mouse models of immunotherapy and vaccination^52–54^. Moreover, convalescent serum from human survivors of CCHFV infection tested positive for antibodies against GP38^52,53^, which we confirmed in this work. In a recent DNA vaccine study, immunized mice failed to be protected when GP38 was not included in the vaccine, and a GP38-only strategy showed a potential protective anti-GP38 IgG threshold in the model^54^. These results indicate a potential protective role in vaccination for GP38. Interestingly, we did not see a strong neutralization response at time of challenge, similar to a previously reported DNA vaccine strategy^16^. Neutralization is not associated with protection in CCHFV mouse models^37,53,55–57^ and circulating neutralizing responses in humans may be low^8,44^, suggesting that protection from CCHFV may be more complex than development of a sterilizing humoral response.

One concern with vaccination is the genetic divergence of CCHFV, and the possibility that a vaccine designed against one clade may not display protection to other clades. To date, only two isolates from distinct clades, Hoti (clade V) and Afghanistan (clade IV)^58^, have been evaluated in the cynomolgus macaque model with each displaying a similar disease course^13,15^. Of note, a cross-clade neutralizing response was observed in subjects infected with either strain^15^. Also, rVSV-CCHFV-GPC was designed from the IbAr10200 strain and showed protection against Hoti in this study. Some vaccines, including rVSV-CCHFV-GPC, have shown protection against heterologous challenges in mice^32,54,59^. Antigenic conservation in Gn/Gc in human cases also suggests that cross-protection is possible^60^. While it will be necessary to develop models for other clades, mouse and NHP data indicate that cross-protection is feasible in CCHFV vaccination.

While a single dose of VSV-CCHFV-GPC did not provide sterilizing immunity, our study demonstrates the efficacy of the vaccine as an anti-CCHFV prophylactic candidate. Viral loads were lower, disease was diminished, and the immune response greater in the vaccinated group. Both humoral- and cellular-driven immunity have been reported in humans suggesting a comprehensive adaptive response may be necessary to maximize protection^61^. However, the goal of this work was to determine the efficacy of a rVSV vaccine to protect from CCHFV disease in a pre-clinical NHP model^62^. Future studies to examine more deeply the cellular and humoral responses elicited by VSV-CCHFV vaccination and natural infection are planned. Recently, work by others have demonstrated the protective potential of NP as a vaccine antigen in a replicating RNA vaccine construct suggesting that a multi-antigen vaccine which couples NP to components of CCHFV M segment may also be worth exploring^63, 64^.

## MATERIALS AND METHODS

### Vaccine design and virus isolates

The ΔGrVSV-CCHFV-GPCΔ vector, herein named VSV-CCHFV-GPC, was previously described^32^. Briefly, construction of the replication competent vector involved the replacement of the VSV glycoprotein coding region with a codon-optimized M coding region of CCHFV strain IbAr10200. The recombinant virus was recovered using an established VSV recovery protocol^62^ and then passaged on baby hamster kidney cells. The final product contained minimal point mutations in the CCHFV M coding region but had a 14 amino acid truncation in the CCHFV Gc C-terminal tail.

A passage 9 stock of Hoti previously used in a macaque model study was generously donated by Dr. Heinz Feldmann (Rocky Mountain Laboratories, Hamilton, MT) ^14^. The stock was passaged on SW-13 cells (ATCC CCL-105) to produce a passage 10 stock as previously described^13^.

### Vaccination and challenge

This study was approved by the University of Texas Medical Branch (UTMB) Institutional Animal Care and Use Committee. The study was performed in the Galveston National Laboratory on the UTMB campus, which is accredited by the Association for Assessment and Accreditation of Laboratory Animal Care International. During procedures, subjects were anesthetized by an IM injection of ketamine or telazol.

Eight healthy adult cynomolgus macaques (*Macaca fascicularis*) of Chinese origin (Worldwide Primates, Inc.) ranging in age from ∼3-6 years old and weighing 3.25-7.50 kg were randomized into two groups. Five macaques (3 males, 2 females) were vaccinated with 2.0×10^7^ PFU of VSV-CCHFV-GPC divided into two equal doses via IM injection of the right and left quadriceps. Three macaques (1 male, 2 females) remained unvaccinated. Thirty-five days post-vaccination, all macaques were challenged with a target dose of 2×10^5^ PFU Hoti, passage 10 (back titer of 4.5×10^5^ PFU) via a single IV injection. Subjects were monitored daily for changes in clinical symptoms such as activity level, feed intake, bleeding, swelling, rashes, respiration, dehydration, and diarrhea. Blood samples, rectal temperature, and weight were collected at pre-planned intervals. At the study endpoint, necropsies were performed and axillary and inguinal lymph nodes, liver with gallbladder, spleen, kidney, adrenal gland, lung, brain (frontal cortex, cerebellum, hippocampus, and stem), cervical spinal cord, pituitary gland, pancreas, urinary bladder, reproductive organs (gonad, uterus, and/ prostate), conjunctiva, and eye tissues were harvested for further analysis.

### Detection of CCHFV

Detection and quantitation of CCHFV vRNA in blood and infectious CCHFV in plasma were performed by RT-qPCR and plaque titration, respectively, as previously described^13^. Briefly, RNA from inactivated blood was extracted, and viral loads were determined using primer-probes that target the NP (5’(6FAM)-AGAAG GGG CTT GAG TGG TT-3’-Dabcyl Quencher 3 (IDT)). Threshold cycle values of CCHFV S genomes were analyzed with CFX Manager Software (BioRad). RNA from CCHFV Hoti was extracted from stock virus and used to generate a genome equivalent standard using Avogadro’s number and the molecular weight of the CCHFV genome. The limit of detection of the assay was 1×10^3^ GEq/ml.

Infectious virus load in serum was detected by infecting SW-13 cells in duplicate with 10-fold serial dilutions of plasma samples. After virus adsorption, cell monolayers were overlaid with 1.25% Avicel RC-581 (FMC BioPolymer) in Eagles minimum essentials medium supplemented with 5% fetal bovine serum (FBS) and 1% penicillin/streptomycin. After four days of incubation at 37°C, 5% CO_2_, the overlay was removed and cells stained with 1% crystal violet in 10% buffered formalin. The limit of detection was 10 PFU/ml.

### Hematology, serum biochemistry, and serum cytokines/chemokines

At sample time points, blood was collected to measure changes in hematology and serum biochemistry parameters. Total counts for white blood cell, red blood cell, thrombocytes, monocytes, neutrophils, eosinophils, basophils, hematocrit, and hemoglobin were measured from whole blood using a Vetscan HM5 Hematology Analyzer (Abaxis). Serum concentrations of albumin, alanine aminotransferase, alkaline phosphatase, amylase, aspartate aminotransferase, blood urea nitrogen, calcium, creatinine, C-reactive protein, gamma glutamyltransferase, glucose, total protein, and uric acid were measured using a Piccolo chemistry analyzer and Biochemistry Panel Plus analyzer discs (Abaxis). Irradiated sera was analyzed for concentrations of circulating cytokines and chemokines, including: L-1β, IL1ra, IL-2, IL-4, IL-5, IL-6, IL-8, IL-10, IL-12/23 (p40), IL-13, IL-15, IL-17, IL-18, IFN-γ, G-CSF, GM-CSF, MCP-1, MIP-1α, MIP-1β, TNF-α, TGF-α, sCD40L, and VEGF. Samples were measured using the Milliplex NHP cytokine magnetic bead panel plex (Millipore Sigma) and read using a Luminex-100 system with data analyzed using Bio-Plex Manager software (Bio-Rad).

### Antibody detection in serum

Antibody response to CCHFV was measured with enzyme-linked immunosorbent assays using macaque or human serum samples. Convalescent patient samples from patients living in Kosovo were donated from CCHFV survivors at the Clinic of Infectious Diseases of University Clinical Center of Kosovo Prishtina, Kosovo), with the approval of the Ethical Committee of the Hospital and University Clinical Service of Kosovo (HUCSK). Samples from CCHFV naïve persons were taken from non-endemic areas under UTMB IRB approval. 96-well plates were coated and incubated at 4°C overnight with a target antigen. Antigens tested include a CCHFV NP, Gc, purified GP38, Gn, Gn/Gc (recombinant proteins provided by Mapp Biopharmaceutical). After blocking with 4% bovine serum albumin in PBS for 2 hours, serial dilutions of serum were added to each well in duplicate with a range from 1:200 to 1:25,600. Serum was incubated for 1 hour before washing and addition of an HRP-conjugated, polyclonal goat anti-monkey IgG detection antibody (Fitzgerald Industries) or HRP conjugated anti-human IgG (Invitrogen). Plates were incubated with secondary antibody for 30 minutes before washing. Signal was detected with a TMB peroxidase substrate system (Sera-Care) and Cytation-5 plate reader (Biotek) at an optical density of 450 nm (OD_450_). Control plates lacking an antigen coating were processed in parallel and utilized to subtract background signal. After background subtraction, samples were normalized to pre-vaccination signal except for control 1. Control 1 lacked pre-challenge samples, so antibody titers were normalized to pre-challenge concentrations. Samples were considered positive if the mean adjusted OD_450_ value of technical duplicates was >0.2. Neutralizing antibody titers in macaque serum were determined by a plaque reduction neutralization test as previously described ^13^. Endpoint titers were assessed by the serum dilution at which a ≥50% reduction in plaques occurs relative to control infections lacking serum.

### Bead-based coagulation assays

Concentrations of circulating coagulopathy-related mediators were determined by flow cytometry using LegendPlex multiplex technology (BioLegend). EDTA plasma levels of each marker were quantified using Human Thrombosis (1:50 dilution) and Human Fibrinolysis (1:10,000) kits (BioLegend). Samples were processed in duplicate following kit recommendations. Following bead staining and wash steps, ∼2100 bead events were acquired using BD FACS Diva software and a FACS Canto II cytometer (BD Biosciences). Raw files were analyzed with cloud-based LEGENDplex™ Data Analysis Software (BioLegend).

### Statistics

All data was graphed and analyzed using GraphPad Prism 8.4.3 software. A Mann-Whitney test was used to determine differences in peak viral loads and time to undetectable titers for both vRNA and infectious virus. Hematological, serum chemical, and coagulopathy biomarkers were normalized to pre-challenge concentrations. No data points were censored in our analysis.

## DATA AVAILABILITY

Complete data sets relating to this work is available upon request from the corresponding author.

## ACKNOWLEDGMENTS

The authors would like to thank the UTMB Animal Resource Center for husbandry support. We would also like to thank Dr. Karla Fenton and Natalie Dobias for assistance with necropsies. Funding was provided in part by the UTMB Department of Microbiology and Immunology to TWG and in part by NIAID/NIH grant number R01AI132246 to TWG. NIAID/NIH grant number UC7AI094660 to UTMB supported BSL-4 operations of the Galveston National Laboratory.

## AUTHOR CONTRIBUTIONS

RWC and TWG conceived and designed the study. RWC, DJD, JBG, TWG carried out *in vivo* activities including vaccination, challenge, and observations. KNA performed PCR and Bioplex assays. VB performed plaque and serum neutralization assays. CW performed Legendplex assays. KM and RO performed the ELISAs. All authors analyzed the data. RWC and KM wrote the paper with additions from AP and TWG. All authors had access to the data and approved the final version of the manuscript.

---

**Supplementary Figure 1:** Changes in clinical parameters in Hoti-infected cynomolgus macaques. Biscuit Count (A) and alanine aminotransferase (ALT, B), and aspartate aminotransferase (AST, C) were measured at sample time points.

## REFERENCES

1 World Health Organization. 2018 Annual review of diseases prioritized under the Research and Development Blueprint. 17 (2018).

2 Zivcec, M., Scholte, F. E., Spiropoulou, C. F., Spengler, J. R. & Bergeron, E. Molecular Insights into Crimean-Congo Hemorrhagic Fever Virus. Viruses 8, 106 (2016). 10.3390/v8040106

3 Hoogstraal, H. The epidemiology of tick-borne Crimean-Congo hemorrhagic fever in Asia, Europe, and Africa. J Med Entomol 15, 307–417 (1979). 10.1093/jmedent/15.4.307

4 Bente, D. A. et al. Crimean-Congo hemorrhagic fever: history, epidemiology, pathogenesis, clinical syndrome and genetic diversity. Antiviral Res 100, 159–189 (2013). 10.1016/j.antiviral.2013.07.006

5 Spengler, J. R. et al. A chronological review of experimental infection studies of the role of wild animals and livestock in the maintenance and transmission of Crimean-Congo hemorrhagic fever virus. Antiviral Res 135, 31–47 (2016). 10.1016/j.antiviral.2016.09.013

6 Ergönül, Ö. Crimean-Congo haemorrhagic fever. The Lancet Infectious Diseases 6, 203–214 (2006). 10.1016/s1473-3099(06)70435-2

7 Spengler, J. R., Bergeron, E. & Spiropoulou, C. F. Crimean-Congo hemorrhagic fever and expansion from endemic regions. Curr Opin Virol 34, 70–78 (2019). 10.1016/j.coviro.2018.12.002

8 Mousavi-Jazi, M., Karlberg, H., Papa, A., Christova, I. & Mirazimi, A. Healthy individuals’ immune response to the Bulgarian Crimean-Congo hemorrhagic fever virus vaccine. Vaccine 30, 6225–6229 (2012). 10.1016/j.vaccine.2012.08.003

9 Bente, D. A. et al. Pathogenesis and immune response of Crimean-Congo hemorrhagic fever virus in a STAT-1 knockout mouse model. J Virol 84, 11089–11100 (2010). 10.1128/JVI.01383-10

10 Bereczky, S. et al. Crimean-Congo hemorrhagic fever virus infection is lethal for adult type I interferon receptor-knockout mice. J Gen Virol 91, 1473–1477 (2010). 10.1099/vir.0.019034-0

11 Zivcec, M. et al. Lethal Crimean-Congo hemorrhagic fever virus infection in interferon alpha/beta receptor knockout mice is associated with high viral loads, proinflammatory responses, and coagulopathy. J Infect Dis 207, 1909–1921 (2013). 10.1093/infdis/jit061

12 Garrison, A. R., Smith, D. R. & Golden, J. W. Animal Models for Crimean-Congo Hemorrhagic Fever Human Disease. Viruses 11 (2019). 10.3390/v11070590

13 Cross, R. W. et al. Crimean-Congo hemorrhagic fever virus strains Hoti and Afghanistan cause viremia and mild clinical disease in cynomolgus monkeys. PLoS Negl Trop Dis 14, e0008637 (2020). 10.1371/journal.pntd.0008637

14 Haddock, E. et al. in Nat Microbiol Vol. 3 556–562 (2018).

15 Smith, D. R. et al. Persistent Crimean-Congo hemorrhagic fever virus infection in the testes and within granulomas of non-human primates with latent tuberculosis. PLoS Pathog 15, e1008050 (2019). 10.1371/journal.ppat.1008050

16 Hawman, D. W. et al. A DNA-based vaccine protects against Crimean-Congo haemorrhagic fever virus disease in a Cynomolgus macaque model. Nat Microbiol (2020). 10.1038/s41564-020-00815-6

17 Daddario-DiCaprio, K. M. et al. Cross-protection against Marburg virus strains by using a live, attenuated recombinant vaccine. J Virol 80, 9659–9666 (2006). 10.1128/JVI.00959-06

18 Falzarano, D. et al. Single immunization with a monovalent vesicular stomatitis virus-based vaccine protects nonhuman primates against heterologous challenge with Bundibugyo ebolavirus. J Infect Dis 204 Suppl 3, S1082–1089 (2011). 10.1093/infdis/jir350

19 Geisbert, T. W. et al. Single-injection vaccine protects nonhuman primates against infection with marburg virus and three species of ebola virus. J Virol 83, 7296–7304 (2009). 10.1128/JVI.00561-09

20 Geisbert, T. W. et al. Vesicular stomatitis virus-based vaccines protect nonhuman primates against aerosol challenge with Ebola and Marburg viruses. Vaccine 26, 6894–6900 (2008). 10.1016/j.vaccine.2008.09.082

21 Mire, C. E. et al. Durability of a vesicular stomatitis virus-based marburg virus vaccine in nonhuman primates. PLoS One 9, e94355 (2014). 10.1371/journal.pone.0094355

22 Mire, C. E. et al. Vesicular stomatitis virus-based vaccines protect nonhuman primates against Bundibugyo ebolavirus. PLoS Negl Trop Dis 7, e2600 (2013). 10.1371/journal.pntd.0002600

23 Marzi, A., Feldmann, F., Geisbert, T. W., Feldmann, H. & Safronetz, D. Vesicular stomatitis virus-based vaccines against Lassa and Ebola viruses. Emerg Infect Dis 21, 305–307 (2015). 10.3201/eid2102.141649

24 Safronetz, D. et al. A recombinant vesicular stomatitis virus-based Lassa fever vaccine protects guinea pigs and macaques against challenge with geographically and genetically distinct Lassa viruses. PLoS Negl Trop Dis 9, e0003736 (2015). 10.1371/journal.pntd.0003736

25 Mire, C. E. et al. Use of Single-Injection Recombinant Vesicular Stomatitis Virus Vaccine to Protect Nonhuman Primates Against Lethal Nipah Virus Disease. Emerg Infect Dis 25, 1144–1152 (2019). 10.3201/eid2506.181620

26 Prescott, J. et al. Single-dose live-attenuated vesicular stomatitis virus-based vaccine protects African green monkeys from Nipah virus disease. Vaccine 33, 2823–2829 (2015). 10.1016/j.vaccine.2015.03.089

27 Marzi, A., et al. EBOLA VACCINE. VSV-EBOV rapidly protects macaques against infection with the 2014/15 Ebola virus outbreak strain. Science 349, 739–742 (2015). 10.1126/science.aab3920

28 Agnandji, S. T. et al. Phase 1 Trials of rVSV Ebola Vaccine in Africa and Europe. N Engl J Med 374, 1647–1660 (2016). 10.1056/NEJMoa1502924

29 Henao-Restrepo, A. M. et al. Efficacy and effectiveness of an rVSV-vectored vaccine in preventing Ebola virus disease: final results from the Guinea ring vaccination, open-label, cluster-randomised trial (Ebola Ca Suffit!). Lancet 389, 505–518 (2017). 10.1016/S0140-6736(16)32621-6

30 Huttner, A. et al. Determinants of antibody persistence across doses and continents after single-dose rVSV-ZEBOV vaccination for Ebola virus disease: an observational cohort study. Lancet Infect Dis 18, 738–748 (2018). 10.1016/S1473-3099(18)30165-8

31 Kennedy, S. B. et al. Phase 2 Placebo-Controlled Trial of Two Vaccines to Prevent Ebola in Liberia. N Engl J Med 377, 1438–1447 (2017). 10.1056/NEJMoa1614067

32 Rodriguez, S. E. et al. Vesicular Stomatitis Virus-Based Vaccine Protects Mice against Crimean-Congo Hemorrhagic Fever. Sci Rep 9, 7755 (2019). 10.1038/s41598-019-44210-6

33 Bodur, H. et al. Evidence of vascular endothelial damage in Crimean-Congo hemorrhagic fever. Int J Infect Dis 14, e704–707 (2010). 10.1016/j.ijid.2010.02.2240

34 Kaya, S. et al. Sequential determination of serum viral titers, virus-specific IgG antibodies, and TNF-alpha, IL-6, IL-10, and IFN-gamma levels in patients with Crimean-Congo hemorrhagic fever. BMC Infect Dis 14, 416 (2014). 10.1186/1471-2334-14-416

35 Ergonul, O., Tuncbilek, S., Baykam, N., Celikbas, A. & Dokuzoguz, B. Evaluation of serum levels of interleukin (IL)-6, IL-10, and tumor necrosis factor-alpha in patients with Crimean-Congo hemorrhagic fever. J Infect Dis 193, 941–944 (2006). 10.1086/500836

36 Swanepoel, R. et al. The clinical pathology of Crimean-Congo hemorrhagic fever. Rev Infect Dis 11 Suppl 4, S794–800 (1989). 10.1093/clinids/11.supplement_4.s794

37 Bertolotti-Ciarlet, A. et al. Cellular localization and antigenic characterization of crimean-congo hemorrhagic fever virus glycoproteins. J Virol 79, 6152–6161 (2005). 10.1128/JVI.79.10.6152-6161.2005

38 Blackburn, N. K., Besselaar, T. G., Shepherd, A. J. & Swanepoel, R. Preparation and use of monoclonal antibodies for identifying Crimean-Congo hemorrhagic fever virus. Am J Trop Med Hyg 37, 392–397 (1987). 10.4269/ajtmh.1987.37.392

39 Cevik, M. A. et al. Viral load as a predictor of outcome in Crimean-Congo hemorrhagic fever. Clin Infect Dis 45, e96–100 (2007). 10.1086/521244

40 Duh, D. et al. Viral load as predictor of Crimean-Congo hemorrhagic fever outcome. Emerg Infect Dis 13, 1769–1772 (2007). 10.3201/eid1311.070222

41 Papa, A. et al. Cytokines as biomarkers of Crimean-Congo hemorrhagic fever. J Med Virol 88, 21–27 (2016). 10.1002/jmv.24312

42 Wolfel, R. et al. Virus detection and monitoring of viral load in Crimean-Congo hemorrhagic fever virus patients. Emerg Infect Dis 13, 1097–1100 (2007). 10.3201/eid1307.070068

43 Burt, F. J., Leman, P. A., Abbott, J. C. & Swanepoel, R. Serodiagnosis of Crimean-Congo haemorrhagic fever. Epidemiol Infect 113, 551–562 (1994). 10.1017/s0950268800068576

44 Shepherd, A. J., Swanepoel, R. & Leman, P. A. Antibody response in Crimean-Congo hemorrhagic fever. Rev Infect Dis 11 Suppl 4, S801–806 (1989). 10.1093/clinids/11.supplement_4.s801

45 Hasanoglu, I. et al. Crucial parameter of the outcome in Crimean Congo hemorrhagic fever: Viral load. J Clin Virol 75, 42–46 (2016). 10.1016/j.jcv.2015.12.006

46 Gurbuz, Y. et al. Evaluation of Prognostic Values of Tissue Plasminogen Activator and Plasminogen Activator Inhibitor-1 in Crimean-Congo Hemorrhagic Fever Patients. Jundishapur J Microbiol 8, e26514 (2015). 10.5812/jjm.26514

47 Medina, L. O. et al. A Recombinant Subunit Based Zika Virus Vaccine Is Efficacious in Non-human Primates. Front Immunol 9, 2464 (2018). 10.3389/fimmu.2018.02464

48 Qin, E. et al. Immunogenicity and protective efficacy in monkeys of purified inactivated Vero-cell SARS vaccine. Vaccine 24, 1028–1034 (2006). 10.1016/j.vaccine.2005.06.038

49 Nakayama, M. et al. Protection against H5N1 highly pathogenic avian and pandemic (H1N1) 2009 influenza virus infection in cynomolgus monkeys by an inactivated H5N1 whole particle vaccine. PLoS One 8, e82740 (2013). 10.1371/journal.pone.0082740

50 Clements, D. E. et al. Development of a recombinant tetravalent dengue virus vaccine: immunogenicity and efficacy studies in mice and monkeys. Vaccine 28, 2705–2715 (2010). 10.1016/j.vaccine.2010.01.022

51 Sanchez, A. J., Vincent, M. J. & Nichol, S. T. Characterization of the glycoproteins of Crimean-Congo hemorrhagic fever virus. J Virol 76, 7263–7275 (2002). 10.1128/jvi.76.14.7263-7275.2002

52 Mishra, A. K. et al. Structure and Characterization of Crimean-Congo Hemorrhagic Fever Virus GP38. J Virol 94 (2020). 10.1128/JVI.02005-19

53 Golden, J. W. et al. GP38-targeting monoclonal antibodies protect adult mice against lethal Crimean-Congo hemorrhagic fever virus infection. Sci Adv 5, eaaw9535 (2019). 10.1126/sciadv.aaw9535

54 Suschak, J. J. et al. A CCHFV DNA vaccine protects against heterologous challenge and establishes GP38 as immunorelevant in mice. NPJ Vaccines 6, 31 (2021). 10.1038/s41541-021-00293-9

55 Garrison, A. R. et al. A DNA vaccine for Crimean-Congo hemorrhagic fever protects against disease and death in two lethal mouse models. PLoS Negl Trop Dis 11, e0005908 (2017). 10.1371/journal.pntd.0005908

56 Hinkula, J. et al. Immunization with DNA Plasmids Coding for Crimean-Congo Hemorrhagic Fever Virus Capsid and Envelope Proteins and/or Virus-Like Particles Induces Protection and Survival in Challenged Mice. J Virol 91 (2017). 10.1128/JVI.02076-16

57 Kortekaas, J. et al. Crimean-Congo Hemorrhagic Fever Virus Subunit Vaccines Induce High Levels of Neutralizing Antibodies But No Protection in STAT1 Knockout Mice. Vector Borne Zoonotic Dis 15, 759–764 (2015). 10.1089/vbz.2015.1855

58 Ramirez de Arellano, E., et al. Phylogenetic Characterization of Crimean-Congo Hemorrhagic Fever Virus, Spain. Emerg Infect Dis 23, 2078–2080 (2017). 10.3201/eid2312.171002

59 Spengler, J. R. et al. Heterologous protection against Crimean-Congo hemorrhagic fever in mice after a single dose of replicon particle vaccine. Antiviral Res 170, 104573 (2019). 10.1016/j.antiviral.2019.104573

60 Ahmed, A. A. et al. Presence of broadly reactive and group-specific neutralizing epitopes on newly described isolates of Crimean-Congo hemorrhagic fever virus. J Gen Virol 86, 3327–3336 (2005). 10.1099/vir.0.81175-0

61 Hawman, D. W. et al. T-Cells and Interferon Gamma Are Necessary for Survival Following Crimean-Congo Hemorrhagic Fever Virus Infection in Mice. Microorganisms 9 (2021). 10.3390/microorganisms9020279

62 Whitt, M. A., Geisbert, T. W. & Mire, C. E. Single-Vector, Single-Injection Recombinant Vesicular Stomatitis Virus Vaccines Against High-Containment Viruses. Methods Mol Biol 1403, 295–311 (2016). 10.1007/978-1-4939-3387-7_16

63 Leventhal, S.S., Shaia, C., Rao, D. et al. Replicating RNA vaccine confers durable immunity against Crimean Congo hemorrhagic fever virus challenge in mice. npj Vaccines 9, 249 (2024). 10.1038/s41541-024-01045-1

64 Hawman, D.W., Leventhal, S.S., Meade-White, K. et al. A replicating RNA vaccine confers protection in a rhesus macaque model of Crimean-Congo hemorrhagic fever. npj Vaccines 9, 86 (2024). 10.1038/s41541-024-00887-z

