## Supplementary figures and images for "A single-dose recombinant rVSVΔG-CCHFV-GPC vaccine protects nonhuman primates against heterologous Crimean-Congo hemorrhagic fever virus challenge"

### SUPPLEMENTAL FIGURE 1

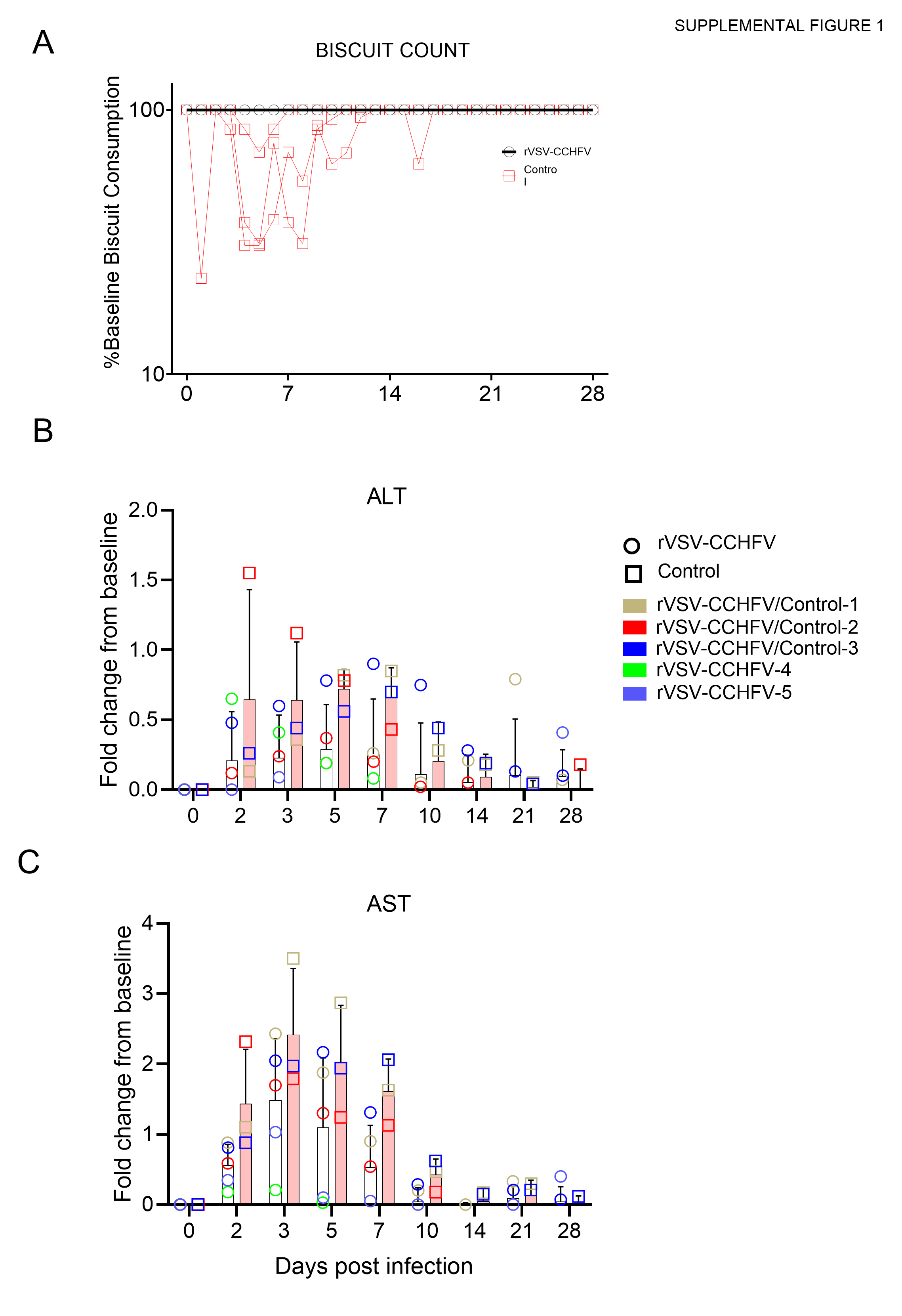
